# Integrated local and systemic communication factors regulate nascent hematopoietic progenitor escape during developmental hematopoiesis

**DOI:** 10.1101/2024.10.21.619535

**Authors:** Carson Shalaby, James Garifallou, Christopher S Thom

**Affiliations:** Division of Neonatology, Children’s Hospital of Philadelphia, Philadelphia, PA, USA; Department of Pediatrics, University of Pennsylvania Perelman School of Medicine, Philadelphia, PA, USA

## Abstract

Mammalian blood cells originate from specialized hemogenic endothelial (HE) cells in major arteries. During the endothelial-to-hematopoietic transition (EHT), nascent hematopoietic stem cells (HSCs) bud from the arterial endothelial wall and enter circulation, destined to colonize the fetal liver before ultimately migrating to the bone marrow. Mechanisms and processes that facilitate EHT and the release of nascent HSCs are incompletely understood, but may involve signaling from neighboring vascular endothelial cells, stromal support cells, circulating preformed hematopoietic cells, and/or systemic factors secreted by distal organs. In this study, we used single cell RNA sequencing analysis from human embryonic cells to identify relevant signaling pathways that support nascent HSC release. In addition to intercellular and secreted signaling modalities that have been previously functionally validated to support EHT and/or developmental hematopoiesis in model systems, we identify several novel modalities with plausible mechanisms to support EHT and HSC release. Our findings paint a portrait of the complex interregulated signals from the local niche, circulating hematopoietic/inflammatory cells, and distal fetal liver that support hematopoiesis.

## INTRODUCTION

The hematopoietic stem cells (HSCs) that support lifelong blood cell production are first produced in major arteries at the onset of cardiac function during embryonic development (Dzierzak and Speck, 2008). Local and systemic signaling mechanisms that support developmental hematopoiesis are incompletely understood. Recent studies have begun to elucidate interactions from non-hematopoietic stromal cells and circulating factors that regulate HSC production (Espín-Palazón et al., 2014; Gonzalez Galofre et al., 2024). Single cell transcriptomics have recently profiled embryonic cells during HSC generation, with a focus on developmental hematopoietic processes (Calvanese et al., 2022). This study was designed to leverage those single cell data to determine signaling mechanisms that impact hematopoietic cell development, focusing on cellular factors both within the hematopoietic niche and across tissues that facilitate HSC formation during embryogenesis.

Hematopoiesis occurs in waves during embryonic development (Dzierzak and Speck, 2008). In all cases, hematopoietic cells are generated from specialized precursors called ‘hemogenic’ endothelial cells (HECs). The first and second waves produce primitive red blood cells, megakaryocytes, and macrophages from yolk sac endothelium. A third wave, which occurs in the dorsal aorta-gonad-mesonephros (AGM) region and other major arteries 4.5 weeks post conception, produces short term multipotent progenitor cells and long term hematopoietic stem cells (Calvanese et al., 2022; Calvanese and Mikkola, 2023). These ‘definitive’ cells transiently colonize the fetal liver from weeks 4.5-6, and ultimately engraft in bone marrow from weeks 8-12 to support lifelong hematopoiesis (Gritz and Hirschi, 2016). Definitive HECs morph into HSCs through an endothelial-to-hematopoietic transition (EHT) and bud off from the aortic endothelial lining. Cells and mechanisms that facilitate HSC ‘escape’ from the endothelial lining are incompletely understood.

Hematopoietic cell-autonomous transcriptional regulatory circuits control HEC specification and fate, including Notch, Wnt, and TGFβ (Ottersbach, 2019; Wu and Hirschi, 2021). In addition, stromal and mesenchymal cells in the arterial niche can support HEC and HSC formation. For example, murine NG2+Runx1+ perivascular smooth muscle cells in close contact to HECs can support hematopoietic development (Wu and Hirschi, 2021; Gonzalez Galofre et al., 2024).

Indeed, explanted AGM tissue can support EHT and HSC formation (Huang et al., 2011; Hadland et al., 2022). Factors in the embryonic circulation also provide signaling input to facilitate HEC specification and HSC formation. Exposure to inflammatory signals, including macrophage-or neutrophil-derived TNFα and via activation of the NFkβ, facilitates HEC specification in zebrafish (Li et al., 2019; Cheng et al., 2023). Augmented inflammatory signals can enhance hematopoiesis from cultured stem cells (Wilken et al., 2024). Less well studied are how signaling post-HEC specification facilitate the escape of nascent HSCs from the arterial endothelial lining.

We hypothesized that resolving local and systemic signals that impact developmental hematopoiesis at single cell resolution would facilitate novel insight into factors that converge to regulate definitive HEC and HSC formation in the AGM, including the initiation and propagation of mechanisms necessary for nascent HSC release. We identified established signaling networks and elucidated novel factors from local and systemic sources, including the fetal liver, that direct developmental hematopoiesis. We additionally identify plausible mechanisms by which inflammatory signaling contributes to EHT, including transcriptional output from signaling events that facilitates cell migration and escape. Harnessing these signaling modalities and mechanisms may boost the quantity or quality of HSCs by better recapitulating the hematopoietic niche in vitro.

## RESULTS

### Single cell analyses identify cells and interactions that support hematopoiesis in the AGM microenvironment

We reprocessed single cell RNA sequencing (scRNAseq) data from human embryonic AGM cells collected at 4.5-6 weeks post-conception (Calvanese et al., 2022). In addition to HECs, HSCs, and mature hematopoietic cells, we identified and annotated stromal and endothelial cell populations (Aran et al., 2019) (**Fig. 1A-B**). We then profiled cell-cell communication within the AGM microenvironment (Jin et al., 2021). We identified 142 unique signaling pathways, composed of 248 ligand-receptor interactions, that target HECs (p<0.05, **Supplemental Tables 1-2**). The calculated ‘probability’ quantifies the relative strength of each interaction compared to all other interactions, providing a way to interpret interaction strengths in the case of extreme P values.

**Figure 1.**
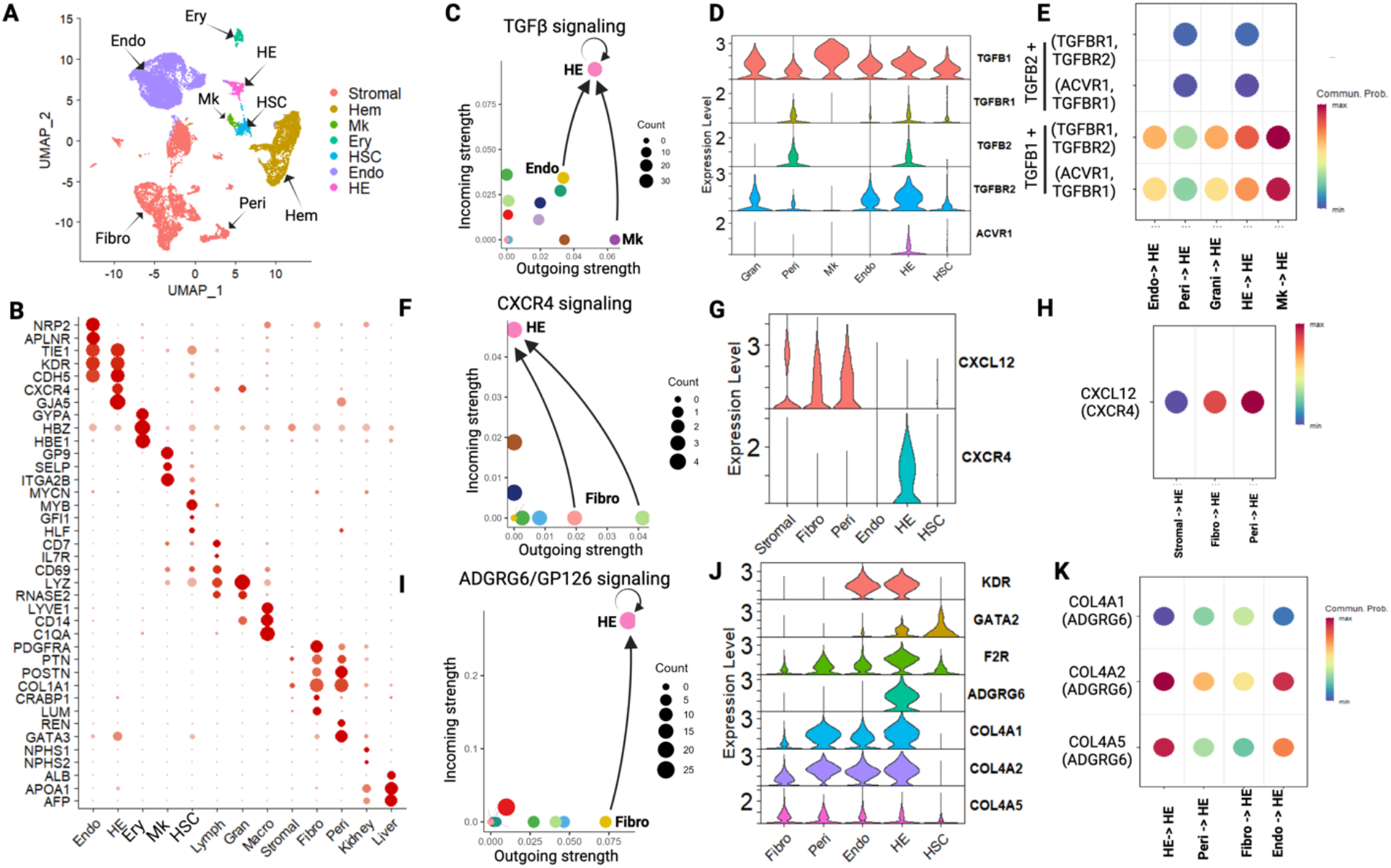
Identification of in vivo signaling mechanisms that regulate hemogenic endothelial (HE) development. A. UMAP depicting populations of cells derived from human embryonic tissue. B. Dot plot supporting annotation of human embryonic cell populations. C. Scatterplot depicting outgoing and incoming TGFβ signaling among analyzed human embryonic cells. HE cells participate in autoregulation and receive signals from ligands produced in megakaryocytes (MKs). D. TGFβ receptor abundance in vascular endothelial and HE cell populations. E.TGFβ ligand-receptor interactions received by HE cells, indicating strongest interactions between TGFβ1 ligand binding to receptor molecules composed of TGFβR1/TGFβR2 or TGFβR1/ACVR1. F. Scatterplot depicting CXCL12-CXCR4 signaling activity in analyzed human embryonic cells. The strongest signaling interaction was between fibroblasts secreting CXCL12 and HE cells expressing CXCR4 receptor. G. CXCR4 receptor expression in vascular endothelial and HE cell populations. H. CXCL12-CXCR4 ligand-receptor interactions received by HE cells indicate that the strongest signaling interactions occur between fibroblast populations and HE cells. I. Scatterplot depicting ADGRG6/GP126 signaling activity in analyzed human embryonic cells. HE cells participate in autoregulation and receive signals from ligands produced from vascular endothelial cells, with substantial ligand production also occurring in pericytes and fibroblasts. J. Collagen ligand and ADGRG6 receptor expression in analyzed cell types. HE cells selectively express ADGRG6 receptor. HE cells, vascular endothelium, pericytes, and fibroblasts express relevant collagen ligands. K. The strongest ADGRG6 communication probabilities support COL4A1/COL4A2 ligand binding to ADGRG6 on HE cells.

Several of these signaling modalities were previously known to support EHT and HSC formation, including TGFβ, Notch, and Wnt (**Fig. 1C** and **Supplemental Fig. 1**). We specifically identified interactions between NOTCH1+JAG1, NOTCH1+DLL4, and TGFb+TGFBR1 that have been functionally shown to facilitate EHT (Butko et al., 2016; Monteiro et al., 2016; Howell et al., 2021). In some cases, these interactions were more prominent in HECs compared to vascular endothelial cells (**Fig. 1C** and **Supplemental Tables 1-2**). However, some ligands produced and/or received at moderate levels by many AGM cell populations, making it difficult to ascertain targeted cell-cell communications. For example, TGFβ receptor expression and communication could potentially occur via ligands produced in several cell types (**Fig. 1D-E**).

### Interactions with stromal cells and extracellular matrix impact EHT

We focused on cases in which HECs were receptive to ligands produced by specific vascular endothelial cells, pericytes, and/or other stromal populations in the AGM environment (**Figure 1F-K**). This approach allowed us to better validate communication modalities via functional evidence and/or targeted bioinformatics strategies. For example, CXCL12 produced by fibroblasts and pericytes is specifically received by CXCR4 on HECs (**Figure 1F-H**). CXCL12-CXCR4 interactions are required for HSC formation (Dignum et al., 2021; Hadland et al., 2022).

We also identified cell interactions between extracellular matrix (ECM) proteins and receptor proteins that were remarkably specific to HECs. For example, the ADGRG6/GP126 receptor which was only appreciably expressed in HECs (**Fig. 1I-J**). ADGRG6 interactions with collagen ligands (COL4A1 or COL4A2) were predominant, as opposed to alternative ligands (PRNP or COL4A5, **Fig. 1J-K**). ADGRG6 receptor interactions with collagen can induce expression of *GATA2* and *VEGFR2/KDR* (Cui et al., 2014) and GATA2 activities can promote EHT (Kang et al., 2018). We noted relevant gene induction downstream of ADGRG6 signaling in HECs (**Fig. 1J**).

Other ECM-Receptor were less specific, although Laminin signaling modalities were enriched in HECs (Aumailley, 2013) (**Supplemental Tables 1-2**). Laminin ligands were produced by several endothelial and stromal populations in the AGM, including HECs themselves. While expression of some Laminin receptors were enriched in HECs vs vascular endothelial cells (e.g., SV2C), the strongest signaling was predicted to occur between Laminins and more widely expressed ITGA9/ITGB1 integrin receptors (**Supplemental Fig. 2**).

These findings confirm and extend the repository of signaling mechanisms that impact HEC development within the AGM niche and lend insight into specific signaling inputs from vascular endothelial and stromal cells that support developmental hematopoiesis. Indeed, some degree of retention in the AGM endothelial lining has been implicated in the ‘arterial education program’ necessary to produce engraftable hematopoietic stem cells (Dignum et al., 2021). However, endothelial retention is ultimately counterproductive to HSC release into circulation. This led us to consider mechanisms responsible for the release of nascent HSCs (or clusters thereof) from the aortic endothelial lining.

### Inflammatory signaling from preformed circulating hematopoietic cells can direct HSC release

Sterile inflammatory signaling directs HEC specification, EHT, and HSC formation in zebrafish (Espín-Palazón et al., 2014), mouse (Li et al., 2019), and cultured human cells (Wilken et al., 2024). We found that TNF produced by macrophages and granulocytes could interact with receptors on human HECs, in alignment with prior findings in model systems (**Figure 2A**). While TNF signaling via TNFR1 was equivalent across all HECs and vascular endothelial cells, only HECs and (to a lesser extent) HSCs expressed TNFR2 (**Fig. 2B-C**). TNFR2 is required for AGM hematopoiesis in the zebrafish (Espín-Palazón et al., 2014) and can initiate different signaling than TNF-TNFR1 binding (MacEwan, 2002).

**Figure 2.**
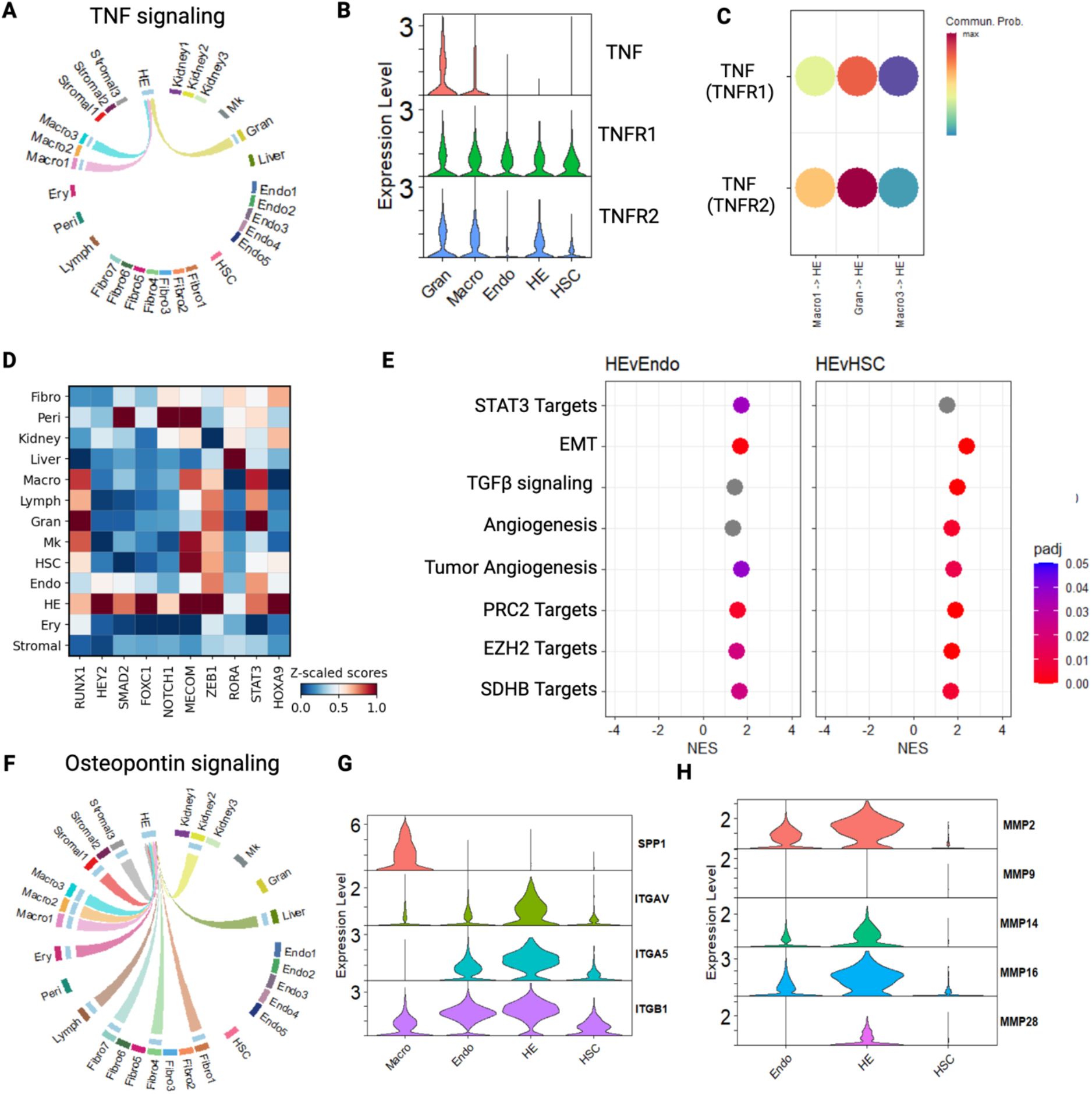
Circulating hematopoietic cells provide inflammatory signaling that enacts downstream transcriptional effects and coordinates nascent HSC escape from arterial endothelium. A. Circle plot depicting TNF signaling input from granulocytes and macrophages received by HE cells. B. TNFR2 expression is enriched in HE cells, whereas TNFR1 expression is consistent across HE cells, vascular endothelium, and HSCs. C. TNF signaling receptor interactions are active across all cell types, although TNF-TNFR2 signaling is selectively active in HE cells. D. Heat plot depicting selected transcription factor-induced gene expression activities in analyzed human embryonic cells. STAT3 activity, a downstream result of TNF signaling, is induced in HE cells vs vascular endothelium. Inflammatory NFkβ is not. Other key transcription factor signaling activities that support developmental hematopoiesis (e.g., RUNX1, NOTCH/HEY2, RORA, HOXA9) are shown for comparison. E. Gene set enrichment analysis (GSEA) showing selected pathway induction in HE cells vs vascular endothelium or HSCs. STAT3 activity is induced in HE cells vs vascular endothelial cells. F. Osteopontin (SPP1) signaling activity in analyzed human embryonic cells. Ligand input from multiple cell types, including inflammatory cells, are selectively received by HE cell receptors. G. Expression of Osteopontin receptor molecules ITGAV and ITGA5 are enriched in HE cells vs vascular endothelium, whereas all endothelial cells express ITGB1. Asterisks represent genes with fold change expression differences >2 (abs[log2FC]>1) with significant differences compared to vascular endothelial cells (Endo, p<0.05). H. Osteopontin signaling can induce expression of matrix metalloproteases (MMPs). MMP2 expression is enriched in HE cells vs vascular endothelium and is known to facilitate escape of nascent HSCs from the arterial endothelial wall during developmental hematopoiesis.

We looked for evidence of downstream activation of TNF-responsive transcription factors within the data. Surprisingly, acute inflammation and other inflammatory pathways were *not* statistically significantly enriched in HEC vs vascular endothelium by gene set enrichment analysis (GSEA). For example, the Inflammatory Response pathway (GOBP) was not statistically enriched in HECs vs vascular endothelium (NES 1.1, p_adj_=0.81). We instead identified alternative transcriptional response to TNFα, including STAT3 target activation in HEC (NES=1.75 vs vascular endothelium, p_adj_=0.036, **Fig. 2D**). In an orthogonal experiment, we used decoupleR to identify transcriptional regulatory networks that were active among sampled embryonic cells. This experiment also identified STAT3 signaling upregulation in HECs vs vascular endothelial cells (**Fig. 2E**). In addition to mediating TNFα-induced inflammatory cytokine production in some circumstances, STAT3 signaling facilitates vascular endothelial permeability and cell migration in the context of tumorigenesis (Azare et al., 2007; Wang et al., 2021). Thus, an inferred effect of TNF-TNFR2 signaling in HECs may be to promote endothelial permeability and EHT/HSC escape.

We further identified regulatory interactions between hematopoietic cells and HECs that have been directly implicated in cell migration and escape from adherent endothelial environments. These signaling pathways included Osteopontin (SPP1/OSN) (Xu et al., 2017; Zhao et al., 2018) (Romacho et al., 2020) (**Fig. 2F**). Osteopontin signaling was mediated by macrophage-derived ligand, although several cell types produce SPP1 (**Fig. 2F-G**). Several integrins can serve as SPP1 receptors, but ITGAV and ITGA5 were notably more highly expressed in HECs vs vascular endothelium (**Fig. 2G**). A key downstream response to Osteopontin signaling is the induction of matrix metalloproteases (MMPs) (Xu et al., 2017; Zhao et al., 2018). In HECs, MMPs facilitate HEC release from the arterial endothelium (Theodore et al., 2017). Indeed, the expression of several MMPs was enriched in HECs vs vascular endothelium (**Fig. 2H**). While MMP2/9 facilitate EHT in mouse, it is likely that analogs more highly expressed in these HECs, such as MMP2, MMP14, and MMP16, mediate this process during human embryonic development.

These experiments begin to paint a systemic portrait of factors that together help initiate developmental hematopoiesis, including stromal support networks and interactions with circulating hematopoietic factors, including a role for inflammatory signaling in the breakdown of the endothelial barrier that would be necessary to release nascent HSCs into circulation.

### Systemic analysis identifies fetal liver-derived support for HSC formation and release

We next extended the scope of potential interactions to include the fetal liver, the site of initial engraftment for HSCs produced in the AGM region (Calvanese and Mikkola, 2023). We hypothesized that the fetal liver might secrete hematopoiesis-inducing factors to indicate its readiness to receive HSCs from the AGM, enabling developmental coordination to promote engraftment in a similar process to bone marrow during later development (Broxmeyer, 2008; Bixel et al., 2017). Indeed, our initial analyses indicated potential liver-derived F2/Thrombin signaling to ADGRG6 on HECs (**Fig. 1F**).

To create a multi-organ single cell data set, we integrated and annotated data from fetal liver and AGM cells, identifying hepatocytes, stellate cells, and activated fibroblast populations among fetal liver cells (MacParland et al., 2018; Giadone et al., 2020) (**Fig. 3A-B**). We then used the full integrated object across all time points to identify 73 unique signaling pathways, composed of 232 ligand-receptor interactions, that targeted HECs (p<0.05, **Supplemental Tables 3-4**).

**Figure 3.**
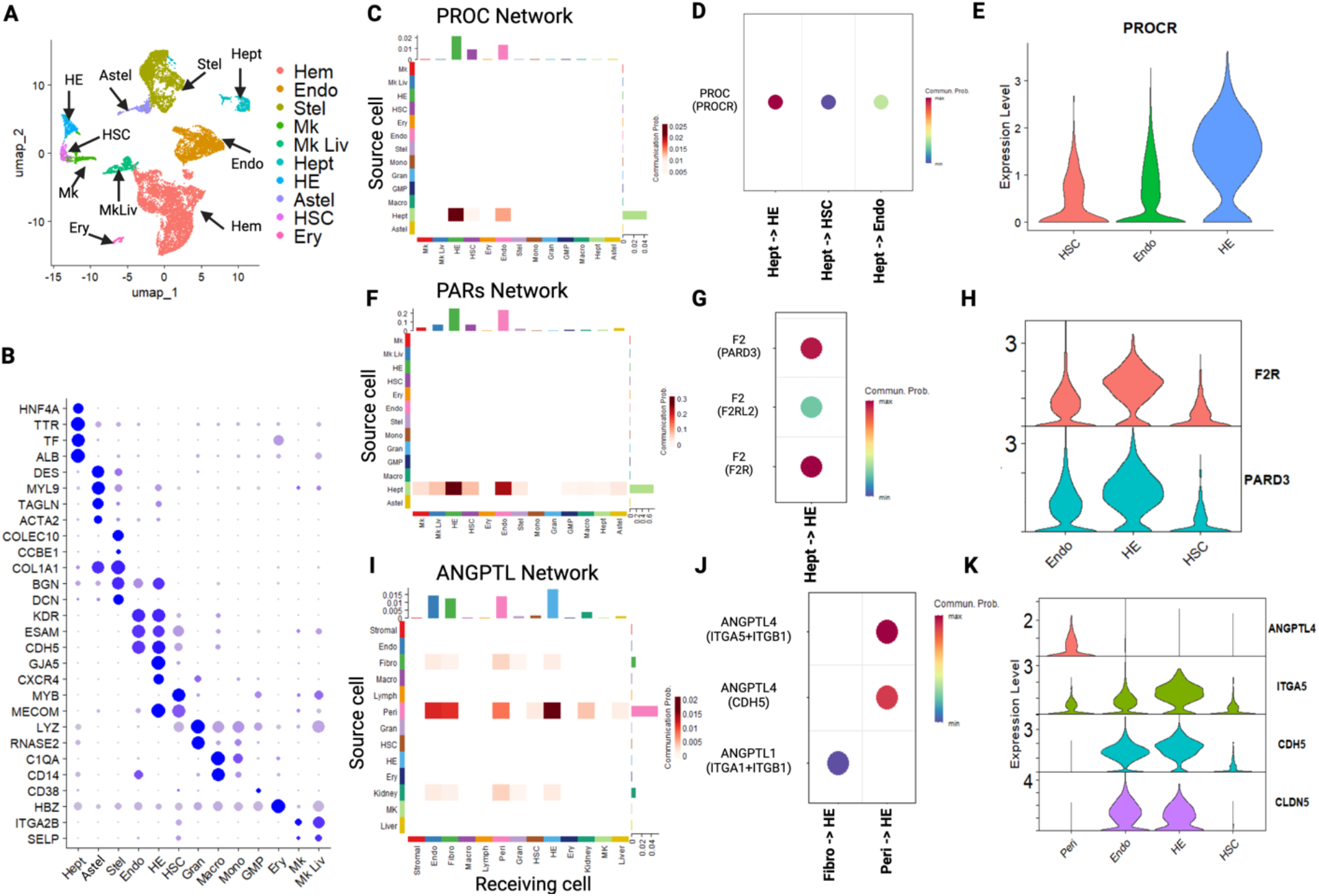
Long range interactions between fetal liver hepatocytes and AGM HE cells regulate developmental hematopoiesis. A. UMAP depicting cell populations from human embryonic tissues, including HE cells, HSCs, and lineage-committed hematopoietic cells from the AGM along with hepatocytes, stellate cells, and activated stellate cells from the fetal liver. B. Dot plot supporting cell type annotations from AGM and fetal liver populations. C. Analysis of Protein C (PROC) signaling activities show selective secretion from fetal liver hepatocytes that is received by HE cells, vascular endothelial cells, and HSCs. D. PROC-PROCR interactions are strongest between fetal liver and HE cells, as compared to vascular endothelium and HSCs. E. PROCR expression is significantly higher in HE cells as compared to vascular endothelium and HSCs. F. Analysis of PAR signaling activities shows that ligands secreted by fetal liver hepatocytes can be received by HE cells and vascular endothelial cells in the AGM. G. Analysis of PAR signaling activities in HE cells shows predominant interactions between fetal liver-derived F2/thrombin ligand and thrombin receptor (F2R) or PARD3 on HE cells. H. Expression of F2R and PARD3 are enriched on HE cells compared to vascular endothelium and HSCs. I. Analysis of Angiopoietin-like (ANGPTL) signaling activities shows that ligands secreted by fetal liver hepatocytes, pericytes, and fibroblasts can be received by HE cells and vascular endothelial cells in the AGM. J. Analysis of ANGPTL signaling activities in HE cells shows predominant interactions between ANGPTL4 and CDH5 and Integrin (ITGA5/ITGB1) receptors on HE cells. K. Expression of ligands and receptors for ANGPTL signaling in AGM cells. ANGPTL4 inhibits tight junctions by binding and sequestering Claudin 5 (CLDN5) and VE-Cadherin (CDH5).

We noted strong Protein C (PROC) and protease-activated receptor (PAR) signaling among fetal liver-HEC interactions (**Figure 3C-H**). In addition to roles in coagulation, PROC and PAR signaling regulates EMT and the regulation of endothelial integrity (Wang et al., 2018; Wojtukiewicz et al., 2019). PROC was exclusively produced by fetal liver hepatocytes, and HECs expressed increased levels of the PROC Receptor (PROC/EPCR) compared with vascular endothelial populations (**Fig. 3C-D**). PROC signaling induces EMT in breast cancer cells (Wang et al., 2018), and EPCR is an established HSC marker that promotes engraftment following irradiation (Balazs et al., 2006; Iwasaki et al., 2010; Lin and Trumpp, 2023). HECs showed 8-fold higher EPCR expression compared with HSCs, and furthermore co-expressed F2R/PAR1 that is necessary for EPCR to mediate PROC-dependent effects (Wojtukiewicz et al., 2019) (**Fig. 3E,H**). PAR signal ligands originating from fetal liver were also relatively specific for HECs and vascular endothelium (**Fig. 3F-G**), with HECs expressing higher levels of F2R and PARD3 than vascular endothelial cells (**Fig. 3G-H**). These signaling interactions may also be involved in breakdown of the AGM endothelial lining, as has been shown in other circumstances (Dragoni et al., 2020).

Finally, we identified novel signaling modalities that may link fetal liver development as a receptive organ to developmental hematopoiesis and EHT. Fetal liver-derived Angiopoietin-like 4 (ANGPTL4) was specifically received by HECs (**Fig. 3I**). ANGPTL4 induces vascular leaks by destroying tight junctions in the endothelial lining (Huang et al., 2011). ANGPTL4 binds to ITGA5B1 and Claudin 5 (CLDN5), which are increased in HECs vs vascular endothelial cells (**Fig. 3J**). In turn, this activates PAK1/Rac, Notch, and Wnt signaling, as well as dissociation of CLDN5 and VE-Cadherin (Cadherin 5, CDH5) at tight junctions (Huang et al., 2011). ANGPTL4 can then bind free CLDN5 and CDH5, preventing tight junction restoration. The liver secretes cleaved ANGPTL4 (cANGPTL4), the most potent version of this gene product that has been linked to cancer metastasis (Huang et al., 2011). In addition to other mechanisms, this represents a novel mechanism by which the fetal liver could plausibly enhance the release of nascent HSCs from the endothelial wall and into circulation (**Fig. 4**).

**Figure 4.**
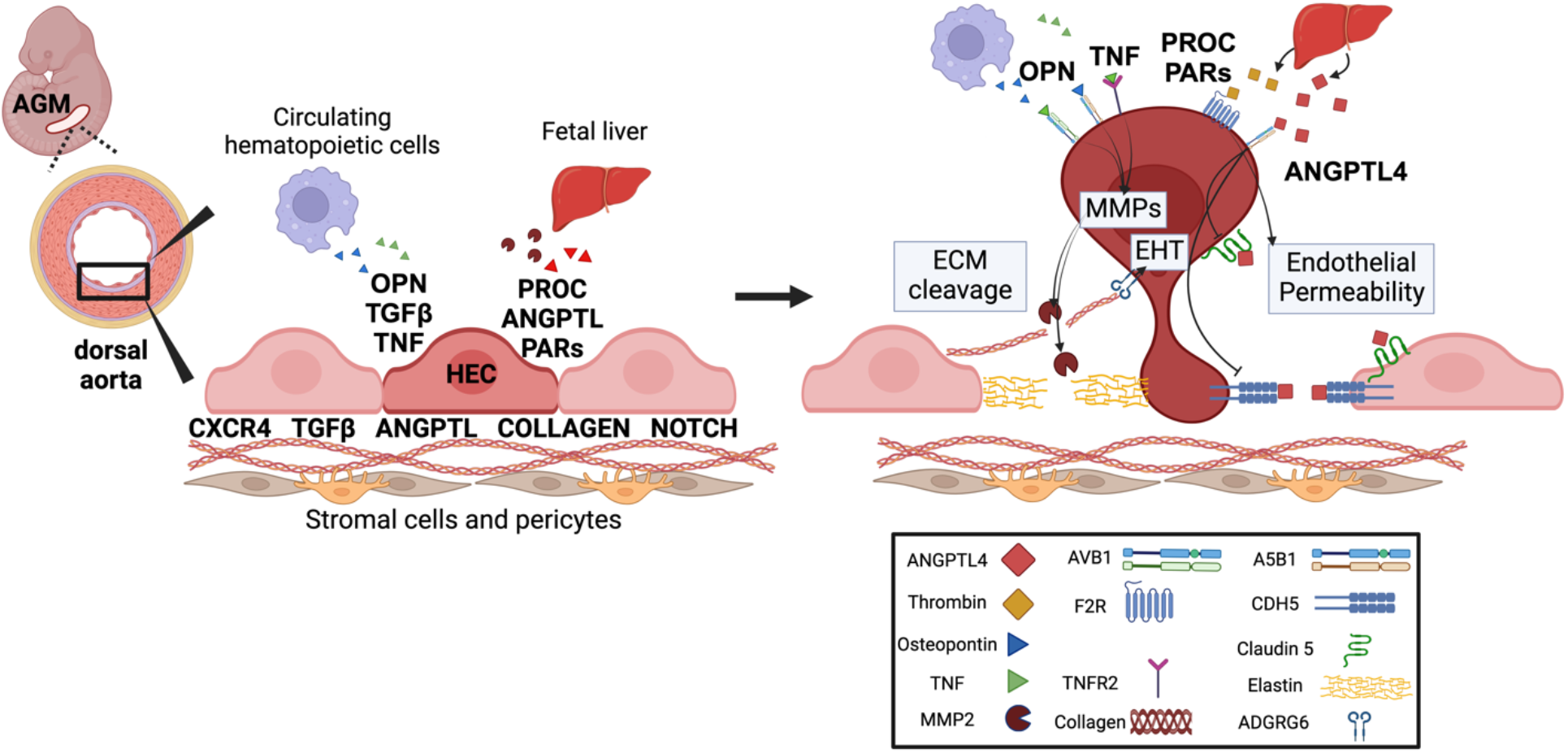
Summary diagram for selected signaling inputs to hemogenic endothelial cells (HEC), with resultant effects on development and ultimately nascent release from the aortic endothelial layer during developmental hematopoiesis. Shown are signals derived from stromal support cells (fibroblasts and pericytes), circulating hematopoietic cells (e.g., macrophages and granulocytes), and fetal liver-derived chemokines that impact the endothelial-to-hematopoietic transition (EHT), endothelial permeability, and extracellular matrix (ECM) cleavage, including matrix metalloproteases (MMPs).

## Discussion

In this study, we identified known and putative signaling inputs that support hematopoiesis via cell autonomous processes and autocrine signaling, as well as supportive interactions with local stroma and circulating inflammatory cells. Interactions within the AGM mirror the systemic factors and microenvironment that regulates hematopoiesis in the bone marrow (Comazzetto et al., 2021), and provide an inclusive view of the factors supporting definitive hematopoiesis in the mammalian embryo. Our findings are consistent with the notion that production of HECs and/or HSCs is a systemic process involving cells outside of the AGM endothelium, including at minimum signals from the fetal liver that may coordinate HSC arrival. Similarly, the fetal bone marrow produces CXCL12 and KITLG to induce migration towards marrow (Miller et al., 2008).

Many of the signaling pathways required for EHT are directly involved with the EMT and associated with cancer progression (Wu and Hirschi, 2021), including mechanisms by which liver signals induce EMT and promote liver metastases (Kim et al., 2006). These parallels are also reflected by gene set enrichment analysis that identified ‘tumor angiogenesis’, but no canonical angiogenesis, as enriched in HECs vs vascular endothelium (**Fig. 2E**). Many of the signaling modalities linked to EHT impact cell morphogenesis and HEC “escape” from the arterial wall, including the dissolution of cell junctions, eliminating apico-basal cell polarity, and adopting a mesenchymal (non-adherent) gene expression pattern. Our findings related to ANGPTL4 and PROC/PAR signaling may reflect discrete mechanisms by which local and systemic signaling might help promote EHT (**Fig. 4**).

Our results further link inflammatory signaling to EHT. Osteopontin (OPN), encoded by the *SPP1* gene, impacts cell migration and cancer metastasis (Xu et al., 2017; Zhao et al., 2018). HECs had increased expression of the OPN-binding integrins AVB1 and A5B1 compared to ECs. ITGAVB1 causes human breast cancer cells to undergo chemotaxis towards OPN gradients, and induces glioma metastasis (Ding et al., 2002; Jan et al., 2010). We also identified downstream targets of OPN signaling, including increased expression of matrix metalloproteases that mediate cleavage of the extracellular matrix during metastasis. MMP2/9 was highly expressed HECs compared to vascular endothelium. MMP2/9 was previously shown to regulate EHT in murine models (Theodore et al., 2017). It is likely that analogous MMPs participate in this process during human development (**Fig. 2H**).

More broadly, our findings align with recent studies showing how hematopoietic cells participate in HEC and HSC formation through production of sterile inflammation. TNF ligand, produced in neutrophils and macrophages and received by HEC, causes downstream NFkβ signaling. This inflammatory signaling mechanism is important for murine (Li et al., 2019), zebrafish (Espín-Palazón et al., 2014; Cheng et al., 2023) and in vitro HSC formation (Wilken et al., 2024). We paradoxically noted downregulation of the TNFR1-induced responses in HECs vs vascular endothelium, likely because of the importance of TNFR2-induced response that is necessary for HEC specification and maintenance (Espín-Palazón et al., 2014).

Our findings indicate that HECs produce basement membrane ligands, like Collagen and Laminins, which participate in autocrine and local signaling interactions. These results may provide novel insight into the biology underlying EHT, specifically the processes by which arterial endothelial cells fill in gaps left following EHT. Extracellular matrix proteins, including Laminins, may help neighboring vascular endothelial cells to migrate and fill in arterial wall space left by migrating HSCs. Arterial wall maintenance is necessary to avoid vascular disintegration and hemorrhage that occurs when HEC specification is enhanced via enforced Runx1+ expression (Yzaguirre et al., 2018). The mechanobiology that supports AGM hematopoiesis, including mechanisms that maintain endothelial integrity during EHT and the release of nascent HSCs, comprises an exciting area for functional interrogation.

This study is based on single cell transcriptomics data and established ligand-receptor databases (Jin et al., 2021). A similar approach has identified important cell communication modalities in physiologic and pathophysiologic microenvironments (Fang et al., 2022). The identification of established control signaling pathways, and links to functionally validated mechanisms, gave us confidence in the presented findings. However, the scope of our experiments was limited to established ligand-receptor databases (Jin et al., 2021). Future spatial transcriptomics and functional validation experiments will further interrogate the importance of specific cell-cell interactions in the developing embryo. Targeted studies may also focus on rare subpopulations that have been functionally linked to developmental hematopoiesis – e.g., a population of RUNX1+ NG2+ perivascular smooth muscle cells recently shown to promote HEC and HSC formation (Gonzalez Galofre et al., 2024).

Defining these active signaling pathways in the complex in vivo hematopoietic niche is important for understanding normal and diseased hematopoiesis, as well as reconstituting hematopoiesis in vitro to produce blood cell-based transfusion products, clinical testing reagents, and other off-the-shelf blood cell-based therapeutics. Prior studies have aimed to recapitulate stromal and mesenchymal cells, which are typically absent from in vitro models. Signaling from pre-formed hematopoietic cells and/or fetal liver may augment in vitro inefficiencies and elucidate new developmental paradigms that influence developmental hematopoiesis.

## Conclusion

This study was designed to test whether in silico methods could detect known cell communications pathways that impact developmental hematopoiesis, forming a basis for the discovery of novel factors that might also temporally impact HEC/HSC formation through local interactions in the AGM and/or distant signaling interactions initiated by other organs or tissues. Through the application of CellChat on a scRNA dataset during definitive hematopoiesis, we identified known and novels signals that impact developmental hematopoiesis. These findings elucidate a more complete portrait of the systemic and environmental signals that direct HEC and HSC formation during human embryonic development, including extracellular matrix interactions and direct contact with neighboring cells in the AGM region, as well as key interactions with circulating hematopoietic cells and long-range secreted signaling interactions with fetal liver.

## Materials and Methods

### Data collection and processing

Single Cell RNA data were obtained from Gene Expression Omnibus (GEO) under accession GSE162950 (Calvanese et al., 2022). We reprocessed combined AGM4.5, AGM5a AGM5b and AGM6 datasets using Harmony to create our completed AGM object which acted as a basis for all future analysis. The AGM+Liver object was created by using Seurat (v5) default tools for combining Seurat objects. Seurat objects representing fetal liver cells at 4.5 5 and 6 weeks were combined with HEC, HSC, and megakaryocyte AGM cells using the merge() function. We restricted analysis to the same study and data sets from Calvanese et al to limit batch/technical effects as able.

Quality control metrics generally aligned with the original manuscript (Calvanese et al., 2022), including 200-8000 features with <10% mitochondrial DNA content. Data were normalized and processed using Seurat (v5.0.1) package functions (NormalizeData, FindVariableFeatures, ScaleData, RunPCA, FindNeighbors, FindClusters, and RunUMAP).

We determined cell and cluster annotations using SingleR (Human Primary Cell Atlas and Bluepoint Encode), CellTypist, ScType, manual curation based on key marker genes, and published annotations (Calvanese et al., 2022). Cells considered to be placental cells were removed. All markers and parameters are found in the supplemental code section and at Github.

### Cell interaction analysis

We used CellChat (v2.1.1) with standard parameters to identify interactions between cells (Jin et al., 2021). We typically examined interactions between individual clusters, rather than cell types, to improve discovery power. All analysis of intercellular signaling was performed at the cluster level (e.g., Endo1, Endo2, Endo3) rather than cell types (e.g., Endo). This was necessary due to heterogeneity among clusters within each cell type. We mandated that >25% cells in a group participated in each ligand-receptor interaction. For simplicity, figures and raw output (**Supplemental Tables 1-4**) are displayed at the cell type level unless otherwise. “TriMean” was used as the argument for the computeCommunProb function. We analyzed interactions with four possible annotations (“Secreted Signaling”, “Cell-Cell Contact”, “ECM-Receptor”, and “Non-Protein Signaling”). Many interaction p-values were extremely low, so we also include calculations for the relative strength of each interaction compared to all other interactions (a value represented as “prob”) in **Supplemental Tables 1-4**. CellChat functions were used to produce graphical output.

### Gene set enrichment analysis

The function “wilcoxauc” from the package “presto” was used to generate a list of marker genes for each cluster of the AGM. The top 100 marker genes of the HE cluster were selected and served as the basis for GSEA analysis. The top marker genes were tested against the 50 Hallmark Gene Sets from the Molecular Signature Database using the package fgsea. This calculated the Net Enrichment Score of each pathway based on the marker genes of the HE.

A similar analysis was performed on an AGM object with only the HE and endothelial clusters to reveal marker genes of the HE relative only to other endothelial cells.

### Transcription factor activity

We assessed signaling pathway output via transcription factor activities using decoupleR. Seurat objects were converted into adata objects using ScCustomize and analyzed in a python3 environment (Ubuntu v22.02). Transcription factor activity inference was performed on the adata object using the CollectTRI database and analyzed using standard parameters.

### Code availability

All data are publicly available from Gene Expression Omnibus (GEO) under accession GSE162950. Analysis code and scripts are posted at https://github.com/okbuddygithub/AGMLIVER and available by request.

## Acknowledgements

This work was supported by the National Institutes of Health (K99 HL156052 to CST).

